# GPR88 localization to primary cilia in neurons is cell-type specific

**DOI:** 10.1101/2025.05.28.656656

**Authors:** Yenni H Li Guan, Brigitte L Kieffer, Mark von Zastrow, Aliza T Ehrlich

**Affiliations:** Department of Psychiatry and Behavioral Sciences, University of California, San Francisco, California, USA; Université de Strasbourg (UNISTRA), INSERM UMR-S 1329, Strasbourg Translational Neuroscience and Psychiatry, Centre de Recherche en Biomédecine de Strasbourg, France

**Keywords:** GPCR, Primary cilia, Orphan receptor, Neuropsychiatry

## Abstract

GPR88 is an orphan G protein-coupled receptor that regulates dopamine neurotransmission and is a target for neuropsychiatric disorders. In addition to the somatic membrane, GPR88 can localize to the primary cilium, a membrane microdomain known for dynamically enriching receptors and signaling molecules. However, the distribution of GPR88 in neuronal primary cilia remains uncharacterized. Here we characterize GPR88 distribution at primary cilia in two brain areas. We show that in the striatum, GPR88 localizes both to somatodendritic and primary cilia compartments on inhibitory GABAergic medium spiny neurons. In contrast, in the somatosensory cortex, GPR88 localizes to somatodendritic and nuclear compartments of excitatory spiny stellate neurons which possess primary cilia that exclude GPR88. Additionally, we found that cilia formation and length were similar between GPR88 knockout and wild-type animals. Together, we provide key evidence for neuronal cell-type specific regulation of GPR88 localization to primary cilia, suggesting neuron subtype specific regulatory mechanisms govern receptor ciliary targeting in the brain.

## Introduction

G protein-coupled receptors (GPCRs) are the largest group of membrane proteins and are involved in critical physiological functions including neuronal function^1,2^. A subset of GPCRs involved in neuromodulation have been found to be targeted to primary cilia^3,4^. Primary cilia are non-motile microtubule-based organelles that protrude off of the cell surface of most cell types in the human body, including neurons throughout the brain^5–7^. In the developing brain, primary cilia are critical locations for Sonic hedgehog (Shh) and Wnt signaling pathways and defects in cilia formation or function can lead to severe disorders known as “ciliopathies”^8,9^. In the mature brain, the function of primary cilia and the GPCRs that enrich it is an emerging area.

Recent work suggests that GPCRs localized to mature primary cilia could be critical for neuromodulation processes including excitatory synaptic regulation^10^ axo-ciliary synaptic regulation of nuclear signaling^11,12^, and energy homeostasis^13,14^. While prototypical ciliary neuromodulatory GPCRs (5HTR6, SSTR3) have been reported to be highly enriched at neuronal primary cilia^15,16^, other ciliary GPCRs lack efficient immunodetection methods to assess ciliary localization in physiological systems. Further, GPCR localization to neuronal primary cilia may be specifically regulated by other proteins or dynamically occur in response to signaling molecules. For example, the dopamine receptor DRD1 is only visible at the primary cilium in brain sections of mice lacking ciliary exit machinery, Bardet-Biedl syndrome proteins^17^, despite being concentrated there in heterologous systems^18^. Additionally, the melanocortin 4 receptor requires an accessory protein to localize to primary cilia that may not be present in heterologous cell systems^19^. Meanwhile the mu opioid receptor is visible at primary cilia in the habenula of wild-type mice but requires an extension of its carboxyl terminus to be transported to primary cilia in heterologous cell culture systems^20^. Thus, to what extent neuromodulatory receptors localize to primary cilia in physiological systems and how GPCR localization to primary cilia is regulated is not clear.

We previously showed that GPR88, an orphan receptor with high interest as a neurotherapeutic target for addiction^21,22^, localizes to primary cilia when overexpressed in ciliated mouse kidney cells and rat striatal neurons^23^ but in physiological systems GPR88 displays time-dependent ciliary enrichment in the nascent period and regional selectivity in the mature brain^24^. The time-dependent nature of GPR88 localization to primary cilia combined with loss of function effects on motor and intellectual ability^25,26^, suggests a role for the receptor in cilia formation or assembly. In contrast the regional restriction of GPR88 to primary cilia on striatal neurons but not cortical neurons suggests that cilia localization of GPR88 may be neuronal context specific.

Here we test the above hypotheses by first systematically identifying neuronal subtypes in the striatum and the somatosensory cortex of GPR88-Venus knock-in mice^24^ and second through detailed quantification of neuronal primary cilia density and length in GPR88-knockout (KO) mice^25^. We find that GPR88 is restricted to SATB2+ excitatory neuronal soma but not cilia in the cortex and DARPP-32+ inhibitory neuronal cilia in the striatum suggesting that GPR88 localization to primary cilia is dependent on the cell subtype. In addition, we find that GPR88 loss of function does not perturb cilia formation or length. These findings suggest that GPR88 loss of function behavioral effects may arise from perturbed GPR88 signaling rather than primary cilia structural or assembly defects. Our results support the concept that ciliary GPCRs require a specific cellular context to be localized to primary cilia.

### Materials and methods Animals

GPR88-Venus knock-in mice were generated at the Institut Clinique de la Souris at Illkirch-Graffenstaden, France. Mice aged 8 to 18 weeks, male and female were bred in-house at the University of California, San Francisco and cared for by Laboratory Animal Resource Center staff. Mice were grouped in cages with free access to food and water and handled according to the guidelines set by the facility. All the procedures were performed according to the guidelines of the Canadian Council of Animal Care and were approved by the Institutional Animal Care and Use Committee (IACUC) at the University of California, San Francisco. Mice were housed with littermates and maintained on a 12-h light/dark cycle with access to food and water.

*GPR88-KO* mouse brains were extracted at the McGill University/Douglas Hospital Research Institute, Montreal Canada and histology procedures were performed at the University of California, San Francisco. *GPR88-KO* mice were generated as previously described^25^.

### Brain tissue preparation

Mice were given CO2 for anesthesia. Wild-type and knock-in, female and male mice were intracardially perfused with 10 mL of PBS (Life Technologies cat# 14040216), followed by 50 mL of 4% PFA/PBS solution (Thermo Fisher Scientific cat#1573590). Heads were cut and brains were extracted into 10 mL 4% PFA and PBS and refrigerated overnight. The whole brains were sunk and were moved into 30% sucrose in PBS for cryoprotection for 48 hours at 4°C. Brains were mounted on deep base molds in dry ice using O.CT. compound (Sakura, VWR) and stored at -80°C until used in immunofluorescence. Brains were sliced in coronal sections at 30 μm on the cryostat (Leica CM1950), sections with the regions of interest (Allen Brain Institute mouse brain atlas, images 44-65) were collected into wells with PBS and kept free floating at 4°C for immunofluorescence procedures.

### Immunohistochemistry and imaging

Sections were permeabilized with PBS-T (0.2% Triton X-100 in PBS) for 10 minutes and then blocked for 1 hour with a blocking buffer 10% normal goat serum (Cell Signaling Technology cat#5425S) in PBS containing 0.3% Triton X-100 (Sigma Aldrich cat#T8787). After, sections were incubated with primary antibodies (**Table 1**) in 3% Normal Goat Serum with PBS overnight at 4°C. Tissue was then washed in PBS-T 3 times for 10 minutes. Secondary antibodies (**Table 2**) were diluted in the blocking buffer and incubated for 1 hour at RT. Sections were then washed in PBS-T for 10 minutes, following a DAPI/PBS wash for 10 minutes, then a final 10 minutes PBS-T wash, slices were mounted onto slides and sealed with coverslips using ProLong gold (Fisher Scientific) and left to dry overnight. All sections were stained with chicken GFP antibody (Novus Biologicals cat#NB1001614) to amplify the GPR88-Venus signal, DAPI (Sigma Aldrich cat#D9542) to stain cell nuclei and rabbit AC3 (abcam cat#ab125093) to label neuronal primary cilia.

**Table 1.** Primary antibodies.

| Antibody | Host species | Dilution | Manufacturer |
| --- | --- | --- | --- |
| Anti-AC3 | Rabbit | 1:1000 | Abcam #ab125093 |
| Anti-ChAT | Goat | 1:500 | Millipore AB144P #3426630 |
| Anti-DARPP-32 | Rabbit | 1:1000 | Cell Signaling Technology #2306S |
| Anti-GFP | Chicken | 1:1000 | Novus Biologicals NB100-1614 |
| Anti-SATB2 | Rabbit | 1:2000 | Abcam #ab34735 |
| Anti-Somatostatin | Mouse | 1:500 | Thermo Fisher Scientific #14-9751-82 |
| Anti-Parvalbumin | Mouse | 1:1000 | Millipore #MAB1572 |

**Table 2.** Secondary antibodies.

| **Secondary Antibody** | **Fluorophore** | **Dilution** | **Manufacturer** |
| --- | --- | --- | --- |
| Donkey anti-chicken | Alexa 488 | 1:2000 | Thermo Fisher Scientific A-78948 |
| Donkey anti-goat | Alexa 647 | 1:2000 | Thermo Fisher Scientific A-21447 |
| Donkey anti-rabbit | Alexa 594 | 1:2000 | Thermo Fisher Scientific A-21207 |
| Goat anti-chicken | Alexa 488 | 1:2000 | Thermo Fisher Scientific A-11039 |
| Goat anti-mouse | Alexa 647 | 1:2000 | Thermo Fisher Scientific A-21235 |
| Goat anti-rabbit | Alexa 594 | 1:2000 | Thermo Fisher Scientific A-11012 |
| Goat anti-rabbit | Alexa 647 | 1:2000 | Thermo Fisher Scientific A-21245 |

For ChAT antibody immunostaining, sections were incubated with 10% normal donkey serum (Jackson ImmunoResearch cat#017-000-121). For SATB2 and DARPP-32, sequential staining was performed as previously reported^20^ (**Figure 2** and **Table S1**). This technique involved staining for the rabbit AC3 antibody first followed by its secondary antibody. The following afternoon a separate same-species antibody was applied for the neuronal marker followed by a secondary antibody. Using this method, the predicted false positives are to detect the cilia marker in both the cilia marker channel and the neuronal subtype marker channel but not vice versa. The GPR88-Venus signal was amplified through species distinct amplification processes and would not be expected to have any false positives. We performed control experiments in which we (1) stained the rabbit raised neuronal subtype markers (SATB2 or DARPP-32) individually from the rabbit raised cilia marker (AC3) and (2) stained only secondary antibodies and omitted the primary antibody. This approach helped us evaluate any non-specific binding of the secondary antibody (see **Figure S2**).

**Figure 1.**
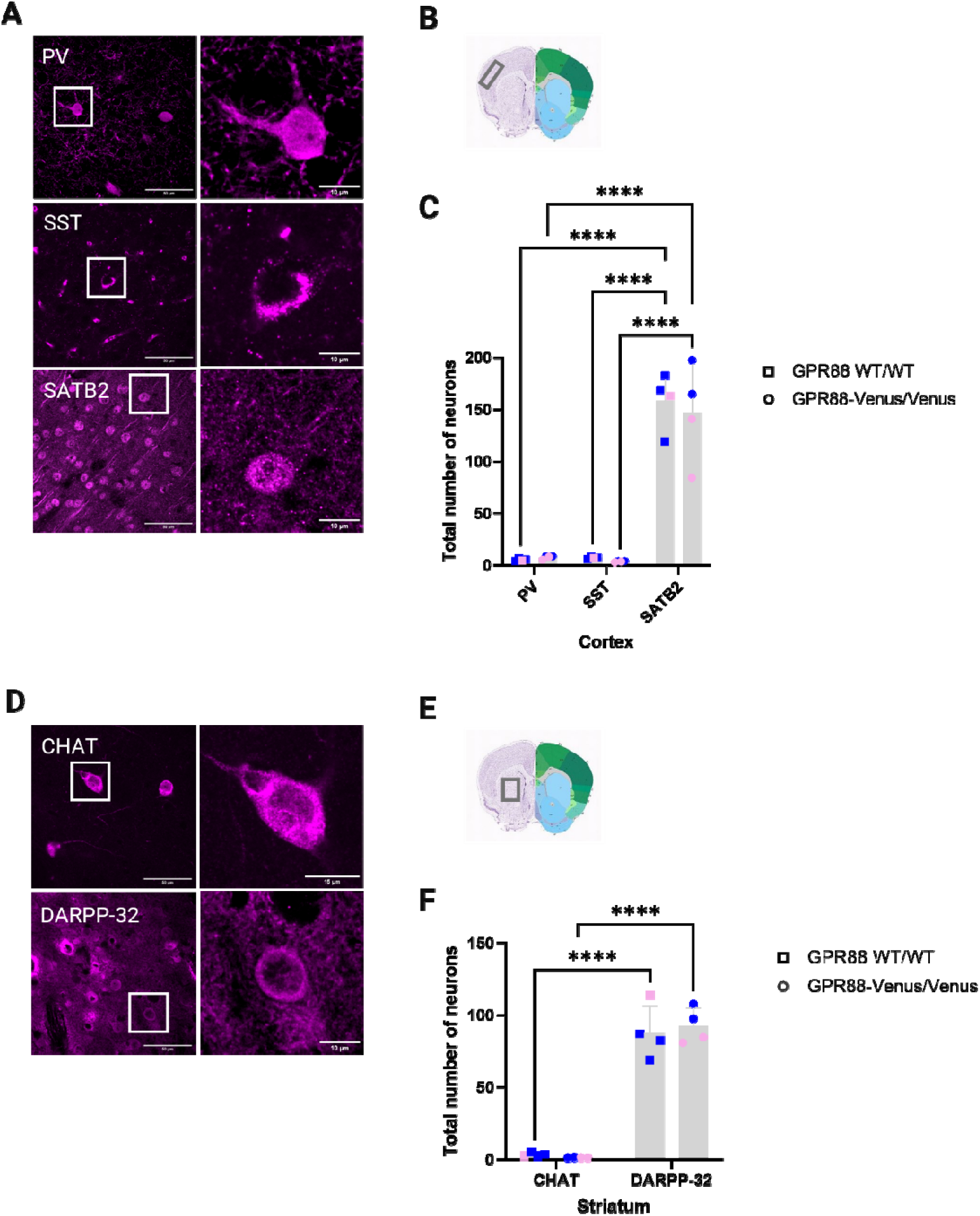
Immunodetection and quantification of cortical and striatal neuronal subtypes. **(A)** Representative confocal images of the SSCtx L4 showing PV, SST and SATB2 expression. **(A**, **D)** Left images show Z-stack projections for each neuronal subtype marker. White boxes indicate representative neurons shown at higher magnification in the images on the right. **(B)** Allen brain atlas image indicates the anatomical location of the images taken in SSCtx L4. **(C)** The bar graph shows the quantification of the total number of neurons expressing each neuronal subtype in the SSCtx L4. The largest population of neurons in the cortex were SATB2+ followed by SST and PV which were observed at similar levels. **(D)** Representative confocal Z-stack projections of the striatum show CHAT and DARPP-32 expression. **(E)** Allen brain atlas image indicates the anatomical location of the images taken in the striatum. **(F)** The bar graph shows the quantification of the total number of neurons expressing each neuronal subtype in the striatum. The largest population of neurons were DARPP-32+ followed by CHAT+ neurons. **(A, D)** Images representing wild-type, adult male mice. **(C, F)** Data from *GPR88^WT/WT^* (n=4; 3 males (blue squares), 1 female (pink squares)) and *GPR88^Venus/Venus^* (n=4; 2 males (blue circles), 2 females (pink circles)). Statistical analysis was performed using two-way ANOVA with Šídák multiple comparison tests, \*\*\*\**P* <0.0001. Data are presented as ± SEM. **(B, E)** Allen Mouse Brain Atlas, https://mouse.brain-map.org/static/atlas.

**Figure 2.**
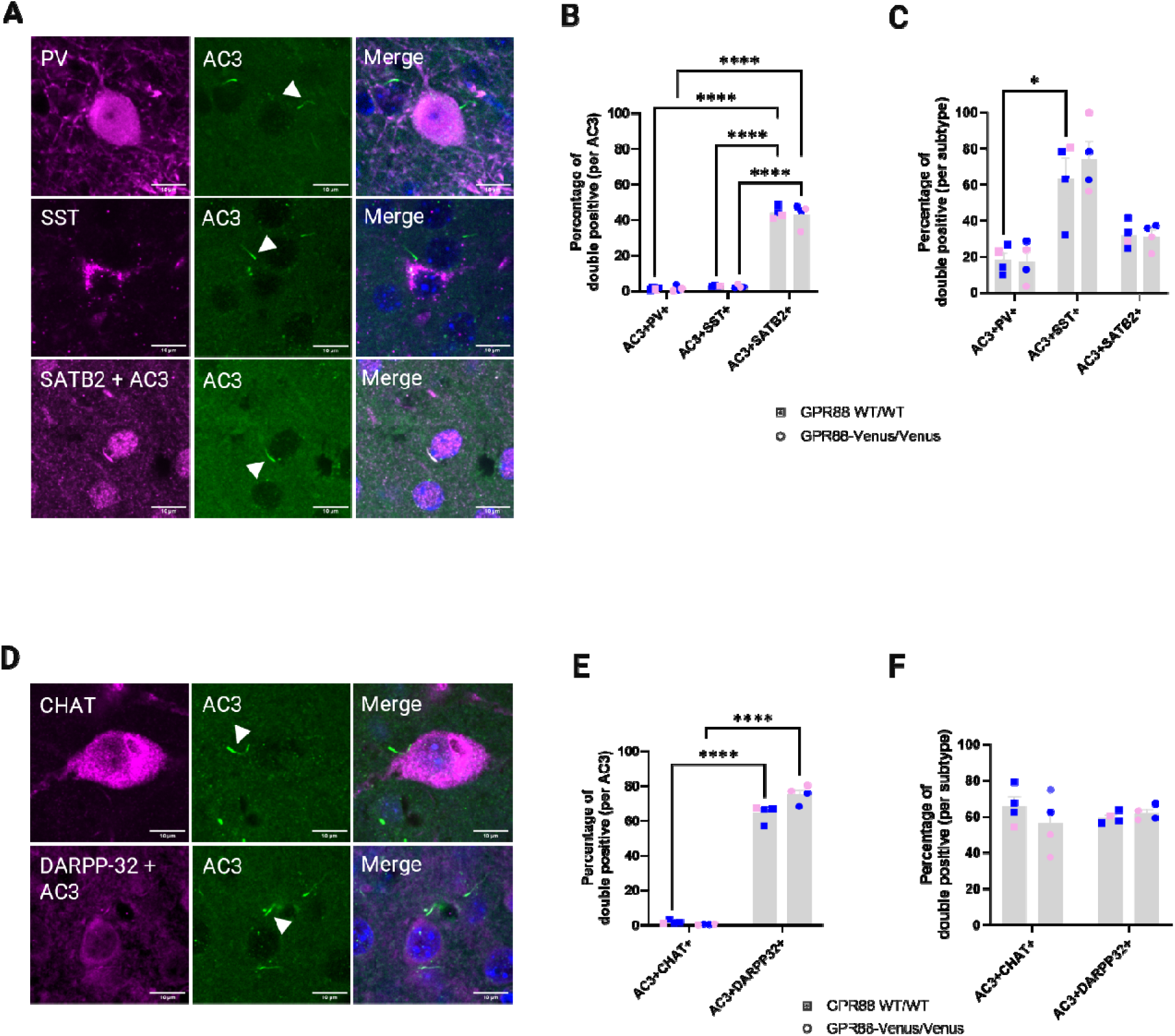
Cilia densities across neuronal subtypes in the somatosensory cortex and striatum. **(A)** Representative images show immunodetection of PV, SST and SATB2 (magenta) in the somatosensory cortex, with AC3 (green) marking primary cilia. Cilia localization is denoted by white arrowheads. AC3 is weakly detected in magenta together with SATB2 and DARPP-32 due to the sequential staining technique (*see method and controls* **Table S1** and **Figure S2)**. **(B)** Quantification of the percentage of double positive per AC3 for cortical subtypes in the cortex shows the distribution of each cell-type in the region.; \*\*\**P* <0.0001. **(C)** Quantification of the percentage of double positive per subtype in the cortex, showing cilia density across PV, SST, and SATB2. \**P* <0.05. **(D)** Representative images showing immunodetection of CHAT and DARPP-32 in the striatum, with AC3 (green) marking cilia. **(E)** Quantification of the percentage of double positive per AC3 for striatal subtypes. \*\*\*\**P* <0.0001. **(F)** Quantification of the percentage of double positive per subtype in the striatum, showing ciliation across CHAT and DARPP-32. Statistical analysis was performed using two-way ANOVA with Šídák multiple comparison tests and data are presented as mean ± SEM. **(A, D)** Images representing wild-type, adult female and male mice. **(B-C, E-F)** Data were obtained from *GPR88^WT/WT^* (n=4) and *GPR88^Venus/Venus^* (n=4) animals. Blue indicates male, pink indicates female.

The immunofluorescence images were acquired using the CSU-W1 high speed widefield microscope (Nikon) with 60x objective lens with oil. For the striatum, four images per section were taken across two sections in the dorsal striatum to assess DARPP-32 and CHAT expression. For SATB2, Parvalbumin and Somatostatin, four images per section were captured in layer 4 of the somatosensory cortex. The Allen Mouse Brain was used as a reference to guide the location and regions of interest (ABI, Atlas: 44-65). Each brain region was imaged as a Z-stack with 0.26 step size.

### Image analysis and quantification

All images were manually quantified using the cell counter plugin in ImageJ. The cell counter and multi-point tools were exploited to mark cells of interest and to keep track of the number of labeled cells in each Z-stack image. Statistical analyses were performed using GraphPad Prism software. Results are presented as the mean ± standard error of the mean (SEM). Graphs were generated using GraphPad Prism.

A challenge in quantification is the potential for high background staining, which can complicate the quantification of positive neurons. During manual counting, we excluded neurons or cilia that were touching borders of the image or did not count objects that were difficult to identify. These criteria were applied consistently across all images to ensure accurate and reproducible measurements.

### Length Measurement and Quantification of Primary Cilia using Artificial Intelligence

Measuring and assessing the structure and length of cilia can be a challenging and time-consuming task. To address this, we implemented an unbiased approach of using AI to accurately measure cilia length and volume. In this method, AI is trained on a set of sample images to identify cilia and segment cilia on the image. Once the training is complete, the AI is applied to a new set of images where it draws a binary mask to identify cilia structures. Following image segmentation, a customized General Analysis (GA) is used to perform various quantitative analyses, including measurements of cilia length and intensity^27^. The results are then exported into a data table, which was easily interpreted and used for further statistical analysis in Graph Pad Prism.

## Results

### Expression and quantification of cortical and striatal neuronal subtypes

To investigate the overarching hypothesis that GPR88 localization to primary cilia is regulated by neuronal subtypes, we first identified and quantified the expression patterns of reported neuronal subtype markers in both somatosensory cortex layer 4 (SSCtx L4) and dorsal striatum (dStr)^28–31^ in GPR88 wild-type (*GPR88^WT/WT^*) and GPR88-Venus (*GPR88^Venus/Venus^*) animals. We identified 3 neuronal subtypes in the cortex and 2 neuronal subtypes in the striatum that could reliably be immunostained with commercial antibodies and were representative of the major neuronal subtypes found in these brain regions^29,30,32–37^. Specifically, we assessed parvalbumin (PV), somatostatin (SST), and special AT-rich sequence-binding protein 2 (SATB2) in the cortex. In the striatum, we looked at choline acetyltransferase (CHAT) and dopamine and cAMP regulated phosphoprotein (DARPP-32).

We first focused on identifying the neuronal subtypes in the SSCtx L4 (**Figure 1A**). Previous reports and our own immunostaining verified that PV and SST were the most abundant inhibitory neuronal markers in SSCtx^29,33^. To label excitatory neurons in SSCtx, SATB2 was selected due to its exclusive expression in this subtype^38^ (**Figure 1A**). We observed that PV and SST neuron’s expression were sparsely distributed across the cortical region while SATB2 expression was denser and more widely distributed. Neuronal quantification was carried out by counting the total number of neurons for each cortical neuronal subtype in the SSCtx L4 (**Figure 1B**). As expected, SATB2+ neurons represented the highest population of neurons (158.5 ± 27.44 for *GPR88^WT/WT^*, and 146.88 ± 47.79 for *GPR88^Venus/Venus^*), followed by PV (5.19 ± 1.21 for *GPR88^Venus/Venus^*) and SST (7.31 ± 1.39 for *GPR88^Venus/Venus^*) (**Figure 1C**). No significant differences in the expression patterns of neuronal subtypes were observed between the GPR88-WT and GPR88-Venus genotypes; however, as expected SATB2+ excitatory neurons were significantly more abundant than PV and SST inhibitory neurons in the SSCtx L4.

Next, we evaluated neuronal subtypes in the striatum. The striatum is predominantly composed of GABAergic medium spiny neurons (MSNs)^31^, and contains sparse distribution of cholinergic interneurons^28^. To distinguish MSNs, we used Anti-DARPP-32 as a specific marker, while Anti-CHAT was used to label cholinergic interneurons (**Figure 1D**). We observed dense widespread labeling of DARPP-32+ neurons and sparse labeling of CHAT+ cholinergic interneurons in the striatum as expected (**Figure 1D-E**). Neuronal quantification further supported that DARPP-32+ neurons were the most abundant (87.94 ± 18.87 for *GPR88^WT/WT^*, and 92.00 ± 11.57 for *GPR88^Venus/Venus^*) and CHAT+ neurons were sparse in comparison (3.25 ± 1.06 for *GPR88^WT/WT^*, and 1.31 ± 0.13 for *GPR88^Venus/Venus^*) (**Figure 1F**). No significant differences were observed between GPR88-WT and GPR88-Venus genotypes, but a significant difference between CHAT+ and DARPP-32+ neuronal counts was observed.

### Primary cilia are found across neuronal subtypes in the somatosensory cortex and striatum

Neuronal primary cilia are microtubule-based organelles that play a key role in a variety of signaling pathways impacting neuronal excitability and behavior^10,39^. In the cortex, cilia are generally shorter making them less apparent as compared to the striatum^40^, where cilia are longer and easier to visualize. We previously reported the observation that GPR88 widely localizes to ARL13B labeled primary cilia in the striatum, whereas cortical neurons do not localize GPR88 to primary cilia^24^. We reasoned that this could be due to incomplete labeling of neuronal primary cilia with ARL13B which predominantly labels glial cells or the lack of primary cilia on cortical GPR88+ neurons. To test this, we compared a known neuronal cilia marker with a wider distribution than ARL13B (**Figure S1**). Adenylyl cyclase 3 (AC3) labeled neuronal primary cilia in the striatum and cortex as expected^40^. ARL13B immunodetection also detected primary cilia in the cortex and striatum but only sporadically. AC3 immunodetection was more frequent compared to ARL13B labeling providing a signal that easily distinguished primary cilia from the neuropil. Therefore, we selected AC3 as the optimal antibody to mark primary cilia on neurons in both brain regions. In cases where the neuronal marker and the cilium antibodies were derived from the same species (anti-SATB2 and anti-DARPP-32), a sequential staining method (see methods and **Table S1**) was employed with standard controls **(Figure S2)** as previously described^20^.

Next, brain sections containing the SSCtx L4 and the striatum from both *GPR88^WT/WT^* and *GPR88^Venus/Venus^* mice were stained by the neuronal subtype marker and the cilia marker (AC3). Primary cilia were clearly evident in confocal images of inhibitory parvalbumin and somatostatin neurons as well as excitatory SATB2+ neurons in the SSCtx L4 (**Figure 2A**). These results show that cortical neurons are ciliated as expected. SATB2 also shows labeling of the AC3+ primary cilium because of the sequential staining method (see methods, **Table S1** and **Figure S2** for more details). Next, we quantified the distribution of ciliated neurons in both regions and calculated the percentage of double positive cells. We quantified two metrics: the ratio of double positive cells (neuronal subtype+ and AC3+) to the cilia marker, AC3, as a metric of overall density across all ciliated cell subtypes or to the neuronal subtype as a metric of cilia density within neuronal subtype. In the cortex (**Figure 2B**), SATB2+ had the highest percentage of double positive per AC3 (43.63 ± 0.6745) compared to other cortical subtypes, PV+ (1.526 ± 0.05486) and SST+ (2.645 ± 0.2614) as expected given that it is known to be the dominant type of neuron in this region^38^. However, when calculating the percentage of ciliated neurons per subtype (**Figure 2C**), SST+ had the highest proportion of ciliated cells (68.92 ± 7.570) compared to the other cortical subtypes, PV+ (17.80 ± 0.7982) and SATB2+ (31.75 ± 0.4493). Statistical analysis revealed no significant differences in the percentage of double positive neurons between *GPR88^WT/WT^*and *GPR88^Venus/Venus^* genotypes in the cortex (**Fig 2B-C**), although significant differences were noted between the cell subtypes (see figure legend). Collectively, this data shows that all cortical neurons possess AC3+ primary cilia with SST+ neurons exhibiting the highest density of ciliated neurons.

To determine cilia density and distribution on striatal neurons we identified cholinergic interneurons with anti-CHAT or GABAergic medium spiny neurons (MSNs) with anti-DARPP-32 and anti-AC3 was used to label neuronal primary cilia (**Figure 2D**). Primary cilia were observed on CHAT and DARPP-32 neurons. Merge images show that cholinergic and GABAergic neurons are ciliated. In the striatum (**Figure 2E**), the percentage of double positive per AC3 was low for CHAT+ (1.297 ± 0.9133) and significantly higher for DARPP-32 (69.92 ± 7.638), the dominant cell type in this region. We also observed a significant difference in the percentage of double positive cells only per AC3 (**Figure 2E**) for DARPP-32 between *GPR88^WT/WT^* and *GPR88^Venus/Venus^*genotypes (genotype F (1, 12) = 6.798, p value = 0.0229) suggesting an increase in DARPP-32+ neurons in the knock-in animals. This finding may be influenced by technical factors or counting considerations, which could have contributed to the observed difference. Interestingly, when we calculated the percentage of ciliated neurons per subtype (**Figure 2F**), the proportion of ciliated CHAT+ and DARPP-32+ were similar between neuronal subtypes and *GPR88^WT/WT^* and *GPR88^Venus/Venus^* genotypes (subtype F (1, 12) = 0.0036, p value = 0.9526, genotype F (1, 12) = 0.5169, p value = 0.4859) suggesting that they exhibit comparable cilia densities.

#### GPR88 is localized to SATB2+ excitatory neurons in the somatosensory cortex and DARPP-32+ GABAergic neurons in the striatum

Next, we asked if the observed regional differences in GPR88 localization to primary cilia are dependent on neuronal subtype. To characterize GPR88 localization to primary cilia in the adult mouse brain, we first determined which neuronal subtype expresses GPR88 using GPR88-Venus mice. We immunostained GFP to enhance fluorescence detection, alongside specific neuronal subtype markers. In the somatosensory cortex (**Figure 3A**), GPR88 was primarily observed on SATB2+ excitatory neurons, with no observed colocalization in inhibitory neurons (PV or SST). The quantitative data further supports these findings. In the cortex (**Figure 3B**), GPR88 expression was only detected in SATB2+ neurons (87.39% ± 1.66) but not in any PV or SST neurons which indicates that GPR88 is specifically expressed in excitatory neurons of the SS Ctx L4. To determine if GPR88 maintains this cell-type specificity at transcript level we turned to the recently published openly available MERFISH transcriptomics dataset from the adult mouse brain^41^. In agreement with our spatial proteomics, *Gpr88* transcripts were only found to co-localize in *Satb2*+ neurons but not *Pvalb* or *Sst* (**Figure 3C-D**) in the SSCtx L4.

**Figure 3.**
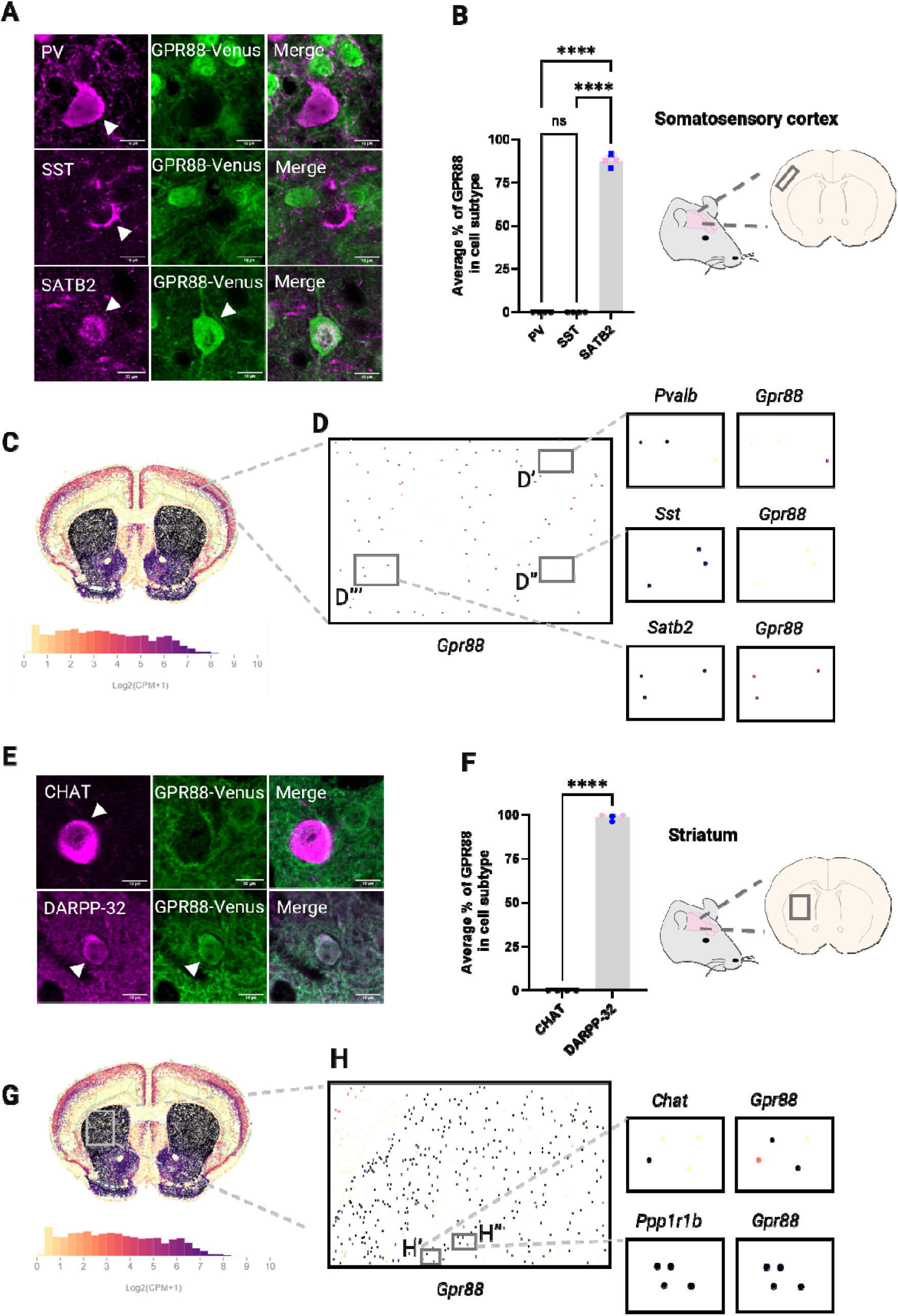
GPR88 is localized to SATB2+ excitatory neurons in the cortex and DARPP-32+ GABAergic neurons in the striatum. **(A-D)** GPR88 distribution in the somatosensory cortex layer 4. **(A-B)** Representative immunofluorescence-stained brain sections that show GPR88-Venus distribution in cortical neuronal subtypes. The GPR88-Venus signal quantified as the average % of GPR88 in a cell subtype is found only in SATB2+ excitatory neurons and is not detected in inhibitory neurons (PV and SST). **(C-D)** Representative spatial transcriptomic images from the Allen Institute MERFISH dataset of adult mouse brain (Allen Brain Cell Atlas (RRID:SCR_024440) https://portal.brain-map.org/atlases-and-data/bkp/abc-atlas) shows *Gpr88* transcripts at whole brain level **(C)** is highest in the striatum (black). At regional level **(D)** in the SSCtx L4 (grey box). *Gpr88* is absent from *Pvalb* **(D’)** and *Sst* **(D’’)** neurons but co-localized with *Satb2* **(D’’’)**. **(E-H)** GPR88 is distributed in DARPP-32+ MSNs in the striatum. **(E-F)** Confocal images of immunofluorescence-stained brain sections and quantification of the average % GPR88 in cell subtype, show GPR88 distribution in DARPP-32 but not CHAT neurons. *Gpr88* at whole brain level **(G)** and **(H)** within the striatum. *Gpr88* transcripts are detected at high level in *Ppp1r1b* (DARPP-32) **(G’’)** MSNs and at lower levels in *ChAT*+ **(G’)** cholinergic interneurons. **(A, E)** Confocal images representing GPR88*^Venus/Venus^* adult female mice. **(B, F)** One-way ANOVA with Tukey’s comparison test; \*\*\*\**P* <0.0001 and values represent the data as ± SEM. Data were obtained from *GPR88^Venus/Venus^*(n=4) animals. Blue indicates male, pink indicates female. **(C-D, G-H)** Transcriptomics expression levels increase in intensity as expression levels increase with yellow (0, not detected) and black as maximal expression. *Gpr88* levels in the striatum are shown as a reference scale (log2(CPM+1)).

In the striatum (**Figure 3E**), GPR88 showed colocalization exclusively with DARPP-32+, MSNs, consistent with previous findings that GPR88 is primarily expressed in these GABAergic neurons^25,42–44^. No colocalization between GPR88 and CHAT was observed, confirming that GPR88 protein is not localized to cholinergic interneurons^43^. The quantitative data in the striatum (**Figure 3F**), show that the average percentage of GPR88 expression in DARPP-32+ neurons was 98.62% ± 0.8798 while CHAT+ was (0 ± 0). This indicates that in the striatum GPR88 receptor protein localizes to MSNs. The transcriptomics were in agreement with DARPP-32 (*Ppp1r1b*) neurons co-localizing with the neurons exhibiting the highest expression of *Gpr88* transcripts (**Figure 3G**) while *ChAT+* neurons contained lower levels of *Gpr88* relative to the MSNs (**Figure 3H**). These results highlight the selective localization of GPR88 protein and transcript to specific neuronal subtypes in both brain regions.

#### GPR88 is only localized to primary cilia in the striatum but not in the somatosensory cortex

To map the distribution of GPR88-Venus to primary cilia in the adult mouse brain, we examined both total and ciliary GPR88 localization in the somatosensory cortex and striatum. Confocal images revealed distinct patterns of GPR88 expression in these regions. In the somatosensory cortex, GPR88 is found on excitatory neurons that are densely clustered together (**Figure 4A**). Confocal images show GPR88-Venus present on soma, neuronal fibers and consistent with previous reports, GPR88-Venus colocalizes with nuclear compartments in cortical neurons^24,43,45,46^. However, even though AC3+ primary cilia was abundant on cortical neurons, no ciliary localization of GPR88-Venus was detected (**Figure 4B, 4E**). This finding confirms the exclusion of GPR88 from cortical primary cilia.

**Figure 4.**
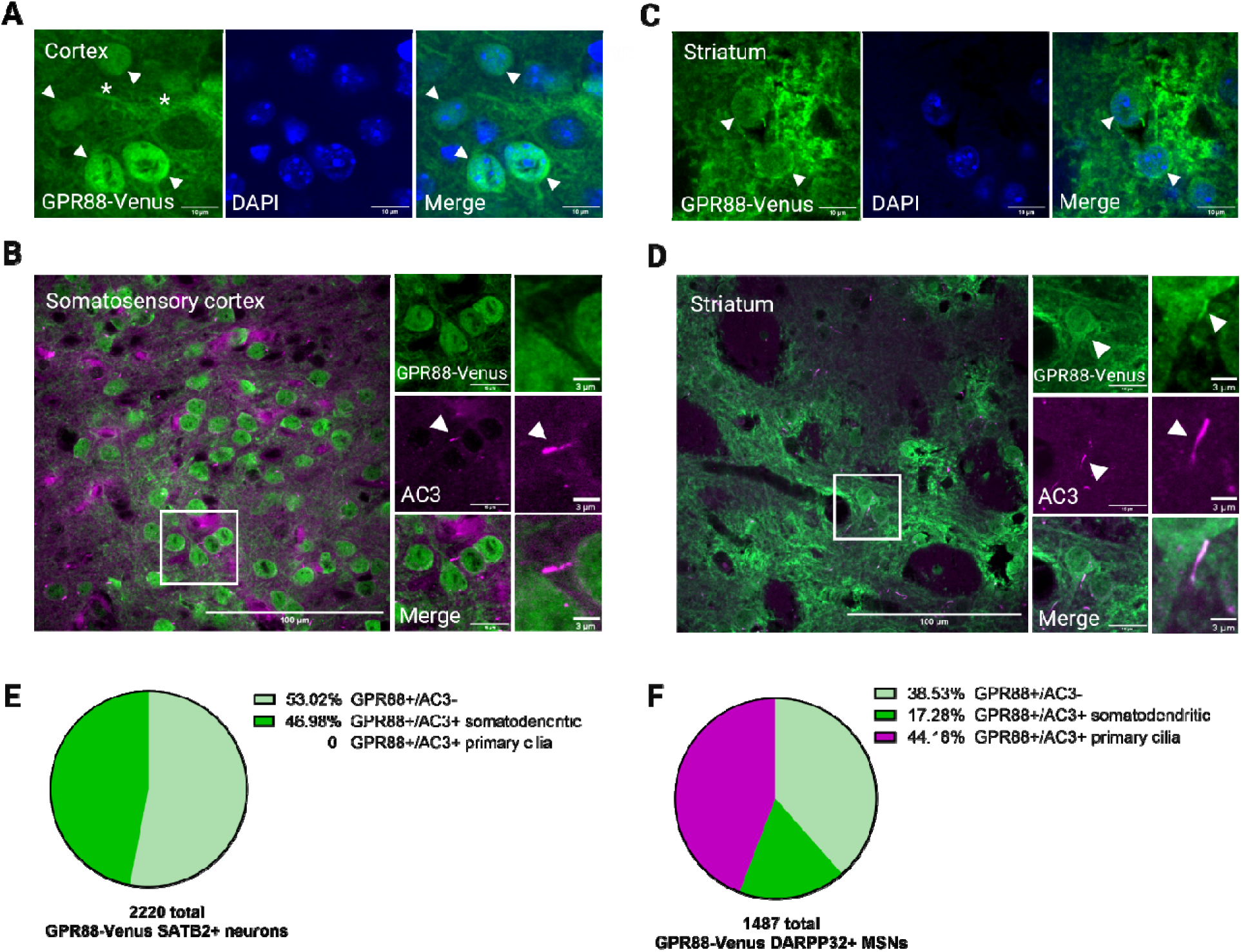
GPR88 is only localized to primary cilia in the striatum, but not in the somatosensory cortex. **(A, C)** Colocalization of GPR88-Venus (green) and DAPI (blue) in cortical and striatal regions. In merged images, arrowheads indicate GPR88-Venus cell bodies. Asterisks indicate neuronal fibers. **(A)** In the cortex, GPR88-Venus colocalizes with DAPI in the nucleus. **(C)** In the striatum, GPR88-Venus is localized to the plasma membrane and primary cilia. Images representing GPR88^Venus/Venus^ **(A)** adult female and **(C)** male mice. **(B)** Confocal images show the overall expression of the GPR88 receptor in SSCtx L4. GPR88-Venus fibers are present, but no ciliary signal is detected; there is no colocalization with AC3. **(E)** The pie chart represents the total count of GPR88-Venus+ neurons in SSCtx L4, with GPR88 distribution in somatodendritic but not primary cilia compartments. **(D)** Confocal images show the overall expression of GPR88 in the striatum. Higher magnification images show GPR88 with the ciliary marker AC3, indicating colocalization to primary cilia. **(F)** The pie chart shows the number of GPR88-Venus+ neurons detected in the striatum and the total count of GPR88-Venus neurons with GPR88 at somatodendritic only or both somatodendritic and primary cilia compartments. **(B, D)** Data were obtained from *GPR88^WT/WT^* (n=4) and *GPR88^Venus/Venus^*(n=4) animals. Blue indicates male, pink indicates female. Representative images are from males.

In the striatum, GPR88 expression is homogeneously distributed throughout the region on inhibitory MSNs (**Figure 4C**). In higher magnification images, GPR88-Venus is found on dense fibers between cell bodies, somatic and primary cilia membranes. Further, colocalization of GPR88-Venus with the cilia marker AC3 confirms that GPR88 is present on primary cilia on mature striatal MSNs (**Figure 4D**). We comprehensively characterized GPR88-Venus+ neurons in the two regions. Remarkably not a single GPR88+/SATB2+ neuron localizes GPR88 receptors to primary cilia in the cortex. In contrast, in the striatum roughly half of all double positive GPR88-Venus+/DARPP-32+ neurons localize GPR88 receptors to primary cilia (**Figure 4F**). This is despite the fact that similar percentages of cortical neurons and striatal neurons were found to possess primary cilia (**Figure 4E, 4F**). These results from the first systematic examination of GPR88 distribution on primary cilia find that GPR88 is localized to primary cilia in a neuronal subtype specific manner, on GABAergic MSNs in the striatum but not on excitatory neurons in the somatosensory cortex.

#### GPR88 does not regulate cilia formation or cilia length

Since cilia are microtubule-based structures critical for axon guidance and neuronal development^47^ and loss of function studies of GPR88 demonstrate extensive remodeling of intracortical and cortico-subcortical networks^48^ we wanted to investigate a role for GPR88 signaling during ciliogenesis. We investigated this by quantifying cilia density and cilia length in GPR88-KO and GPR88-WT animals using an unbiased approach with trained artificial intelligence (AI) methods^27^. Brain sections from GPR88-WT and GPR88-KO animals containing the two regions, somatosensory cortex and striatum, were sectioned and immunostained for AC3 to stain primary cilia and DAPI to stain nuclei (**Figure 5A-D**). Higher magnification images show that cortical primary cilia appear shorter (**Figure 5A**) than the longer striatal cilia (**Figure 5C**). These differences reached significance when cilia length was quantified (see **Figure S3**). Comparison of cilia densities (AC3/DAPI) revealed no significant differences between GPR88-WT and GPR88-KO animals in the cortex (**Figure 5E**) and striatum (**Figure 5G**). Additionally, there was no difference between males and females in the striatum. Next, we compared cilia length in the two regions. Measuring the length of thousands of primary cilia (7495 cilia) revealed no significant differences in cilia length between GPR88-WT and GPR88-KO animals in the cortex (**Figure 5F**) and striatum (**Figure 5H**). Further there was no difference between males and females in the striatum. These results suggest that deletion of GPR88 does not affect cilia formation or assembly in the striatum or cortex.

**Figure 5.**
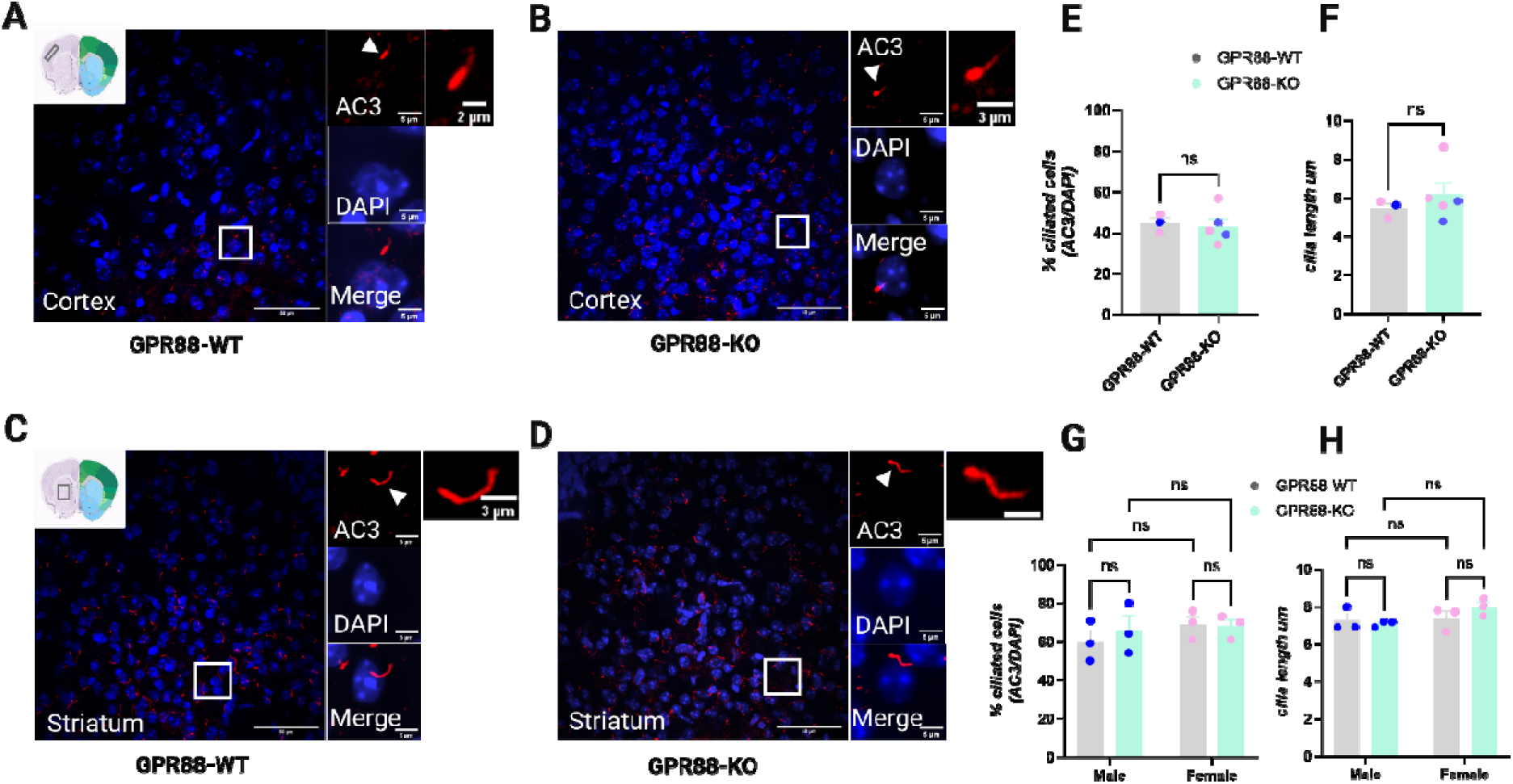
Cilia length and density are unaffected by GPR88 depletion. Representative confocal images showing GPR88 expression in **(A, C)** GPR88-WT and **(B, D)** GPR88-KO mice. **(E)** Bar graph displays the percentage of ciliated cells in the cortex by genotypes, which is measured by AC3/DAPI. **(F)** Graph shows the average cilia length in µm for both genotypes. **(G)** Bar graph shows the percentage of ciliated cells in the striatum by gender, measured by AC3/DAPI. **(H)** Average of cilia length is shown in µm for both genders. Statistical analysis was performed using a two-way ANOVA, Tukey’s test. No significant differences were observed between the genders or genotypes in either cilia density or length. GPR88-WT n=3 (cortex), n=6 (striatum), GPR88-KO mice n=5 (cortex), n=6 (striatum). Blue indicates male, pink indicates female.

#### Variation of GPR88 distribution in the striatal cilia

Since ciliary GPCRs are signaling molecules, they are known to alter their distribution in primary cilia dynamically ^49^. In MSNs, GPR88 was observed in primary cilia of many but not all GPR88+ neurons (**Figure 4F**) suggesting that GPR88 may be variably translocated to the cilium. Therefore, we assessed the distribution of GPR88-Venus within AC3 marked cilia at steady-state. In the striatum, we noticed two types of primary cilia distribution patterns of GPR88-Venus (**Figure 6A, C**). Higher magnification images confirmed the partial colocalization of GPR88-Venus and AC3, with GPR88-Venus expression predominantly concentrated at one end of the cilium (**Figure 6A-B**), in contrast to the longer AC3 labeling along the full-length of the axoneme of the cilium. This pattern of partial GPR88 localization was observed in approximately 40% of the GPR88-Venus brain sections. Interestingly, we also observed approximately 60% of the time a more uniform distribution of GPR88 along the full-length of the cilium (both proximal and distal) (**Figure 6C-D**). The higher magnification images further confirmed that GPR88-Venus and AC3 colocalized evenly along the cilium. This variation in GPR88 distribution across striatal cilia may reflect different states of GPR88 activity, suggesting that GPR88 localization could be dynamic and activity-dependent in this region.

**Figure 6.**
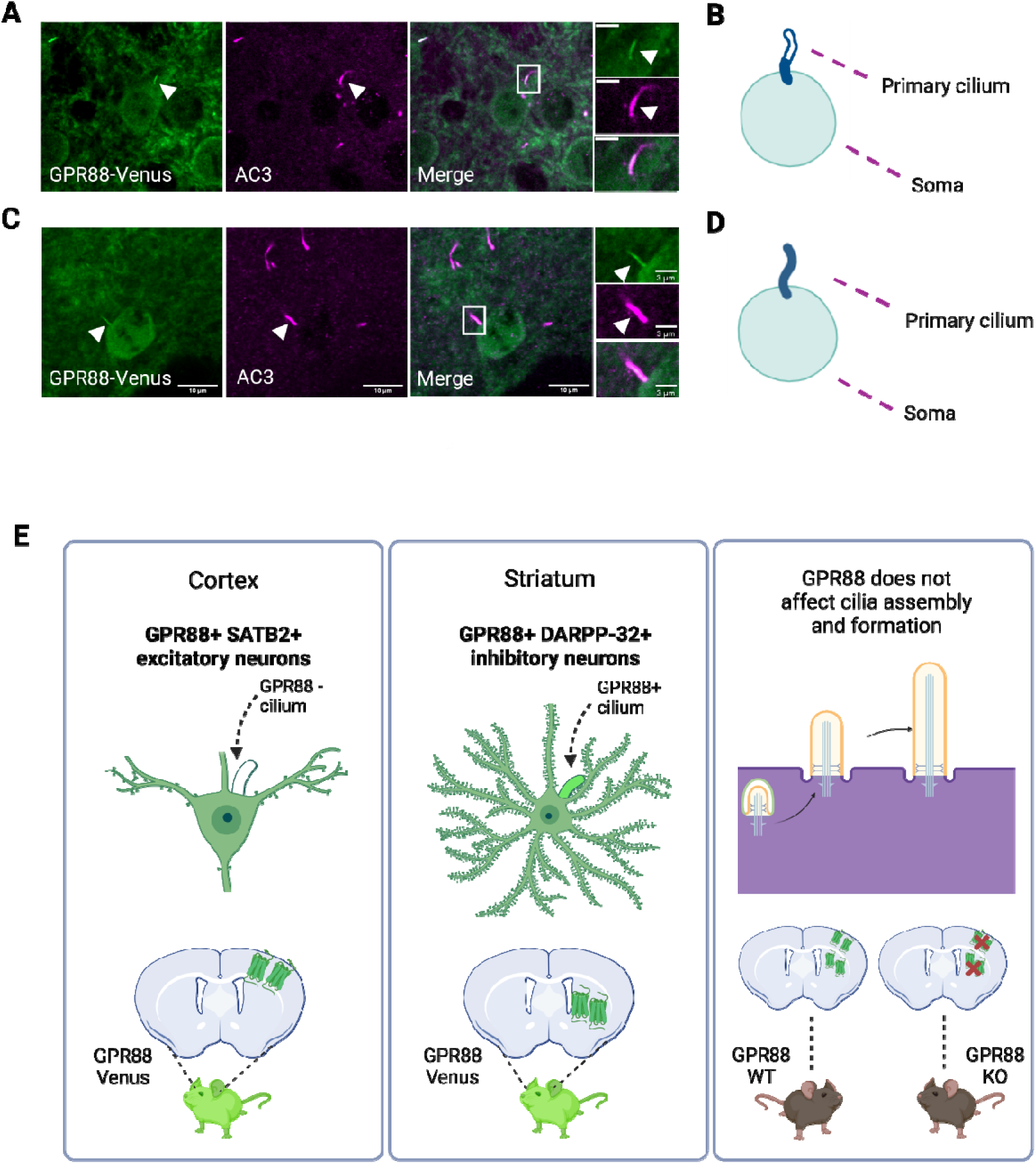
Distribution of GPR88-Venus is variable within striatal primary cilia. **(A, C)** Immunofluorescence staining in GPR88*^Venus/Venus^*mice in the striatum. GPR88-Venus (green) reveals the different distribution patterns of ciliary GPR88, while AC3 (magenta) labels primary cilia. White boxes indicating regions in higher magnification images show the colocalization of GPR88-Venus with AC3 varies along the cilium. **(B)** Scheme depicting partial labeling of GPR88 accumulated at one end of the cilium near the soma. **(D)** Schematic representation illustrates uniform GPR88 distribution along the complete AC3 labeled length of the primary cilium. Representative images from adult **(A)** female and **(C)** male GPR88-Venus mice, n=4. White arrowheads indicate cilia. **(E)** Summary scheme shows (i) GPR88 is exclusively found in excitatory SATB2+ neurons in the SSCtx L4 but excluded from the primary cilium (left panel), (ii) GPR88 localizes to primary cilia exclusively in DARPP-32+ medium spiny neurons in the striatum (middle panel) and (iii) GPR88 function is not required for primary cilia formation and assembly (right panel). Scheme generated with BioRender.com.

## Discussion

GPR88 is an orphan receptor highly expressed in the striatum where it modulates dopamine and thus is a potential target for neuropsychiatric disorders such as ADHD, substance abuse, and Parkinson’s disease^21,44,50–52^. Recent advancements in single cell transcriptomics reveals that neurons express dozens of GPCRs rather than a single GPCR^53^. These findings suggest that determining the spatial organization of receptors is integral to understanding GPCR biology^53,54^. To improve drug targeting strategies for GPR88, a better understanding of its spatial organization is needed. Here, we show that GPR88 localization to primary cilia differs across neuronal subtypes and that GPR88 is not functionally involved in cilia formation. Specifically, we show that GPR88 (1) localizes to primary cilia on inhibitory DARPP-32+ GABAergic MSNs in the striatum; (2) is excluded from primary cilia on GPR88+ excitatory SATB2+ neurons in the cortex; and (3) is not required for cilia formation and assembly which proceed normally in GPR88-KO animals (**Figure 6E**). This work establishes neuronal cell-type specific regulation of GPR88 localization to primary cilia. Ciliary targeting of GPCRs is well known to be receptor-specific, as defined by some GPCRs localizing to cilia and others being excluded. The present results provide a clear example of cell type-specificity in ciliary targeting among distinct neuronal cell types, and, to our knowledge, are the first to show cell type-specific ciliary targeting of any GPCR *in vivo*. Such cell type-specificity adds to a growing body of evidence supporting the general hypothesis that the primary cilium is functionally important for neuromodulation, and it opens the door to future studies testing this hypothesis specifically for GPR88

### GPR88 localizes to excitatory SATB2+ spiny stellate cells in the cortex

Our investigation into the hypothesis that GPR88 localization to primary cilia might be regulated by neuronal subtypes yielded clear findings at cellular and subcellular levels. At the cellular level, we found that GPR88 is restricted to excitatory SATB2+ spiny stellate cells. These cells are primarily found in the SSCtx L4 and receive input from the thalamus participating in cortical functions including sensory processing and information transfer between cortical areas^55,56^. A functional implication for these neurons in sensorimotor connectivity is supported by our previous results from brain-wide fMRI, where loss of GPR88 was shown to have significant alterations in sensorimotor cortical connectivity^57^ and from our behavioral investigation of sensory processing deficits in GPR88-KO animals which exhibit reduced tactile responses^24^. In contrast with another observation^45^, GPR88 was not observed on any of the inhibitory cortical interneurons we examined, including those expressing PV and SST. This discrepancy may arise from study differences either in the region of focus in the cortex or GPR88 detection methods. Notably, the MERFISH transcript dataset in the Allen Brain Cell Atlas also supports our finding that GPR88 localizes specifically to SATB2+ neurons in the SSCtx L4 (**Figure 3**). Future studies that specifically manipulate GPR88/SATB2 cortical neurons will clarify the role for GPR88 neuromodulation in sensory processing.

### GPR88 localizes to DARPP-32 medium spiny neurons in the striatum

The striatum is a region known for high expression of GPR88^42–44^, particularly in MSNs. However, the functional role of GPR88 at the primary cilium of MSNs remains poorly understood. Our results show that GPR88 is localized specifically to DARPP-32+ MSNs and not to cholinergic interneurons (CHAT+) (**Figure 3C**) as previously reported by others^43^. These findings support a functional role for GPR88 MSNs in mediating direct and indirect dopamine pathways and behavior in the striatum as supported extensively by GPR88-KO studies^25,50,58–61^.

### Absence of GPR88 in cortical primary cilia

At the subcellular level, we found that GPR88 localizes to the cytoplasm and nucleoplasm as originally reported by Massart and colleagues^45^ but is excluded from the primary cilium on cortical SATB2+ neurons. Electron microscopy data also showed unconventional GPR88 localization to the nuclear compartment together with biochemical interaction between GPR88 and nuclear proteins suggesting a role for GPR88 in modulating chromatin accessibility in cortical neurons^46^. Our finding of GPR88 exclusion from primary cilia in the cortex has not been mechanistically determined and could arise from several factors. GPR88 localization to cilia may be regulated by developmental control as it was shown to change levels of enrichment at cilia in the striatum as the animal matures^24^. Future studies using GPR88-Venus animals to map GPR88 expansion to cilia across developmental time points in the cortex may help reveal temporal cilia dynamics of GPR88 in the cortex. Additionally, distinct molecular weight species of GPR88 could account for our findings, however similar full-length GPR88 forms have been detected in cortical and striatal lysates^46^. *Cis*-acting ciliary targeting sequences could also be a factor and are generally found on the receptor carboxyl terminus (C-terminus)^62^, the same receptor region that receives post-translational modifications (PTM) that impact localization^63^. It is conceivable then that GPR88 might require a PTM on the C-terminus to be targeted to the primary cilium in cortical neurons. Notably, GPR88 localization to the cilium has not been detected anywhere in the brain by a C-terminus recognizing antibody detection method suggesting it does not recognize ciliary GPR88^43,45^. Recently, *trans*-acting factors have also been identified in transport to cilia. For example, MC4R, was shown to require an accessory protein, MRAP2, to localize to primary cilia in IMCD3 cells^64^ and Rhodopsin has long been shown to require a set of specific accessory proteins to localize to the cilia-like outer segment^65^. In non-neuronal cells, GPR88 transport to the primary cilium has been shown to require a ciliary adaptor protein, TULP3^66^ and it is tempting to speculate that TULP3 or a yet to be identified ciliary adaptor protein may be specifically lacking in cortical neurons. Future work is needed to identify the *cis* and *trans*-acting mechanisms of GPR88 delivery to primary cilia. Collectively, these findings suggest GPR88 localization to primary cilia may be regulated by both temporal and cell-specific mechanisms.

### GPR88 localization to primary cilia on MSNs

Additionally, our findings confirm that ciliary GPR88 is localized exclusively to the striatum and not the cortex^24^. Plasma membrane localization of GPR88 in the striatum, along with its presence on primary cilia, provides a novel perspective on the potential functional significance of GPR88 in MSNs, particularly in cAMP signaling. The primary cilium is known to be a restricted compartment that amplifies cAMP signaling^67^. The enrichment of GPR88 at the primary cilium suggests that the Gαi–GPCR inhibits ciliary cAMP signaling in MSNs. In the paraventricular nucleus of the hypothalamus, overexpression of a mutationally active GPR88 was shown to decrease the activity of ciliary localized Gαs-coupled MC4R^14^. It is interesting to speculate that GPR88 could play a similar role in MSNs, perhaps insulating and buffering the cAMP signal generated by dopamine receptors as has been shown for D1 and GPR88 in IMCD3 cells^18,23^. GPCR localization to primary cilia is generally thought to be dynamic^68^. Our study also sheds light on the dynamic role for GPR88. In the striatum, we observed that ciliary GPR88 is not uniformly distributed along the cilium, with a subset of cilia displaying accumulation of GPR88 at one end of the cilium (**Figure 6A**). This partial localization of GPR88 within cilia suggests that the receptor may be involved in localized signaling mechanisms at the ciliary base or tip in MSNs. Alternatively, it may suggest that GPR88 is dynamically trafficked in and out of the cilium depending on signaling similar to smoothened which depends on Hedgehog signaling^4,69,70^. Additionally, since DARPP-32 labels all MSNs, and we only identified partial localization of GPR88 to primary cilia it would be worthwhile for future studies to determine whether ciliary GPR88 is preferentially localized to D1 or D2 MSNs, or whether it is present in both populations.

Recent studies in striatal neurons and other cell lines have demonstrated that the local receptor environment is critical for determining the cAMP/PKA response of a GPCR^71,72^. The local receptor environment at the primary cilium through modulating cAMP/PKA signaling likely affects neuromodulation either through transcriptional regulation^67^ or regulation of synaptic neurotransmission^6,7,11,73^ regulating complex biological processes such as energy homeostasis^19^ or cognition^74^. Future work precisely manipulating GPR88 localization at MSN primary cilia and measuring signaling effects will clarify the role ciliary GPR88 localization plays in dopamine regulated behavior. Collectively, these results suggest that GPR88 is selectively targeted to primary cilia in the adult striatum and its dynamic localization points to a potential role for ciliary GPR88 signaling in the regulation of striatal neuron function in the mature brain, particularly in the context of neuropsychiatric disorders that involve altered dopaminergic signaling.

## Acknowledgement

We thank the M. von Zastrow lab for the insightful comments and feedback. We thank the UCSF Center for Advanced Light Microscopy (K. Herrington, S. Kim) for providing services and assistance for technical help. We thank the Julius lab and the Rubenstein lab for the use of the cryostat. We thank the Core facility ICS, Michel Bouvier and Brigitte Kieffer for the creation of the GPR88-Venus mouse.

This work was supported by National Institutes of Mental Health grants R01MH12012 (MVZ and BLK), and K01MH123757 (ATE).

## Authors Contributions

Y Li Guan: Data curation, formal analysis, investigation, visualization, methodology and writing original draft, review and editing.

BL Kieffer: Resources, funding acquisition, manuscript review and editing

M von Zastrow: Supervision, funding acquisition, manuscript review and editing

AT Ehrlich: Conceptualization, data curation, resources, supervision, validation, methodology, project administration and writing original draft, review and editing, funding acquisition.

## Conflict of Interest Statement

The authors declare that they have no conflict of interest.

## Notes

### Competing Interest Statement

The authors have declared no competing interest.

