## Supplemental Figures and Tables for "GPR88 localization to primary cilia in neurons is cell-type specific"

**Supplemental Materials:**

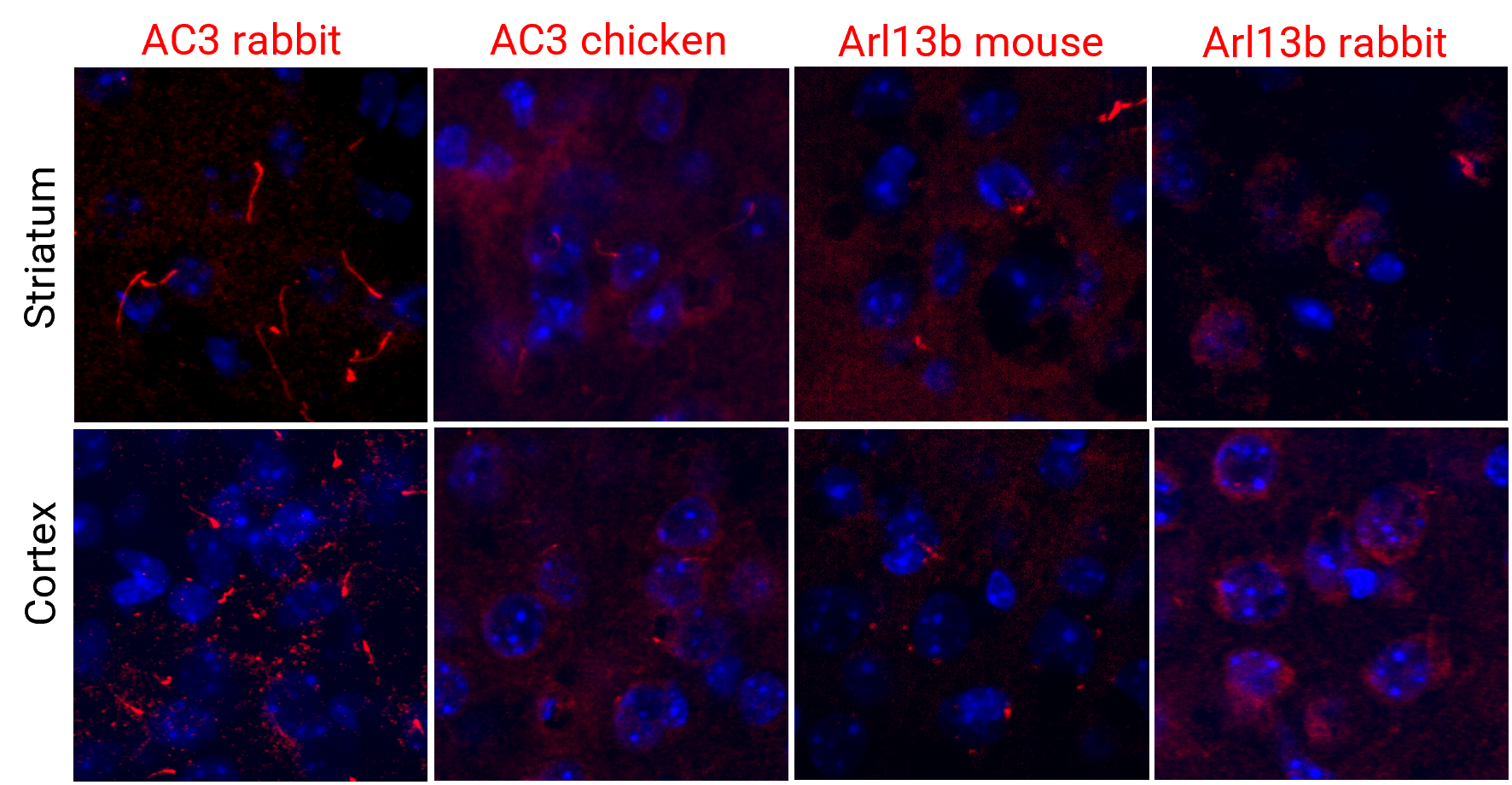

**Figure S1. Comparison of AC3 and ARL13B cilia markers used in immunofluorescence staining.** Expression of primary cilia markers in WT mouse brain. Red represents a cilia marker on each image (rabbit, chicken AC3 and mouse, rabbit ARL13B). All nuclei were stained with DAPI (blue).

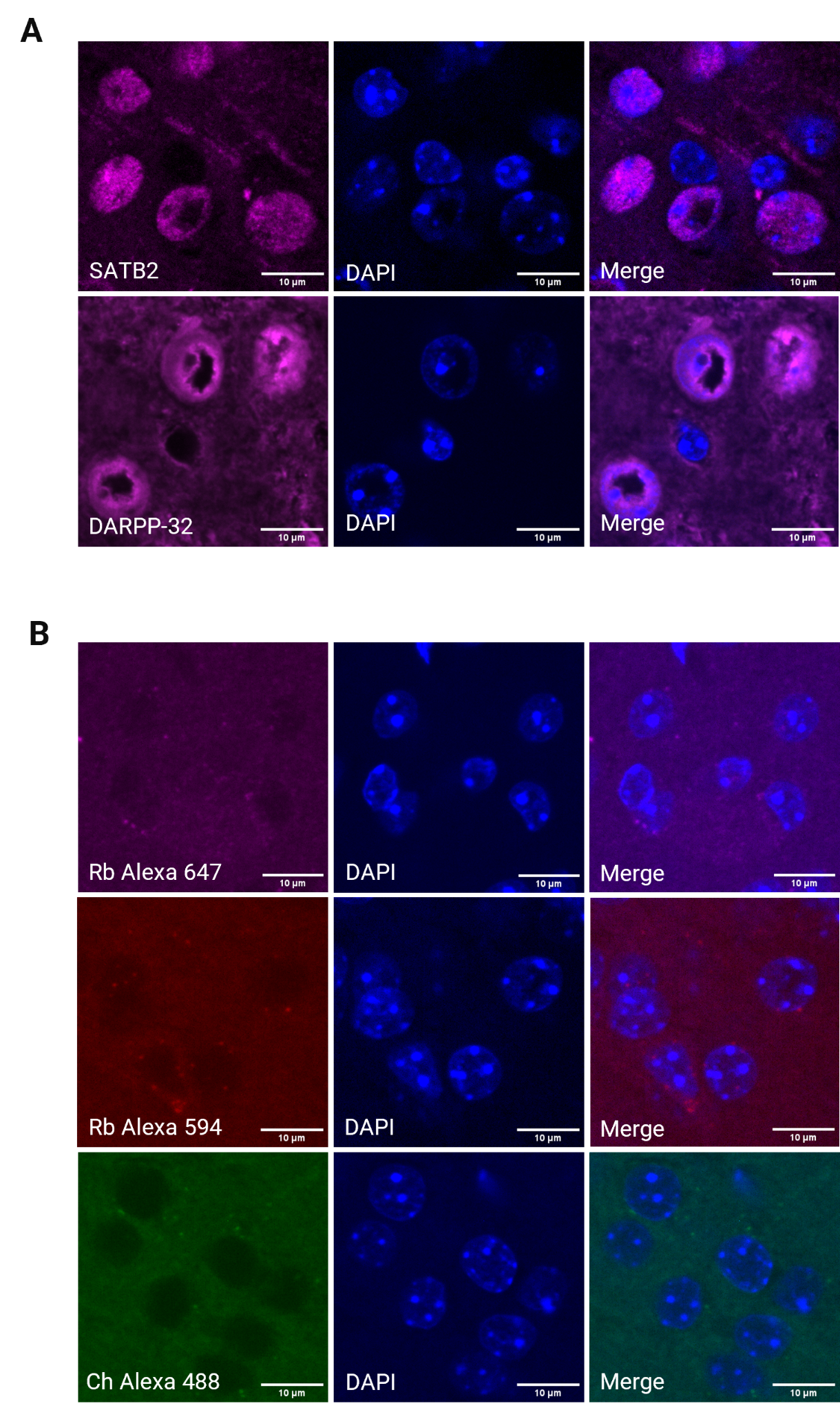

**Figure S2. Immunofluorescence control images of SATB2, DARPP-32 and negative control. (A)** Control images of individual staining of SATB2 and DARPP-32 to show that there are no cilia (AC3). **(B)** Negative control staining was performed without the primary antibody, demonstrating no specific binding by the secondary antibody (Alexa Fluor 647 Goat Anti-rabbit, Alexa Fluor 594 Goat Anti-rabbit and Alexa Fluor 488 Goat Anti-chicken). Images representing wild-type, **(A)** adult male and **(B)** female mice.

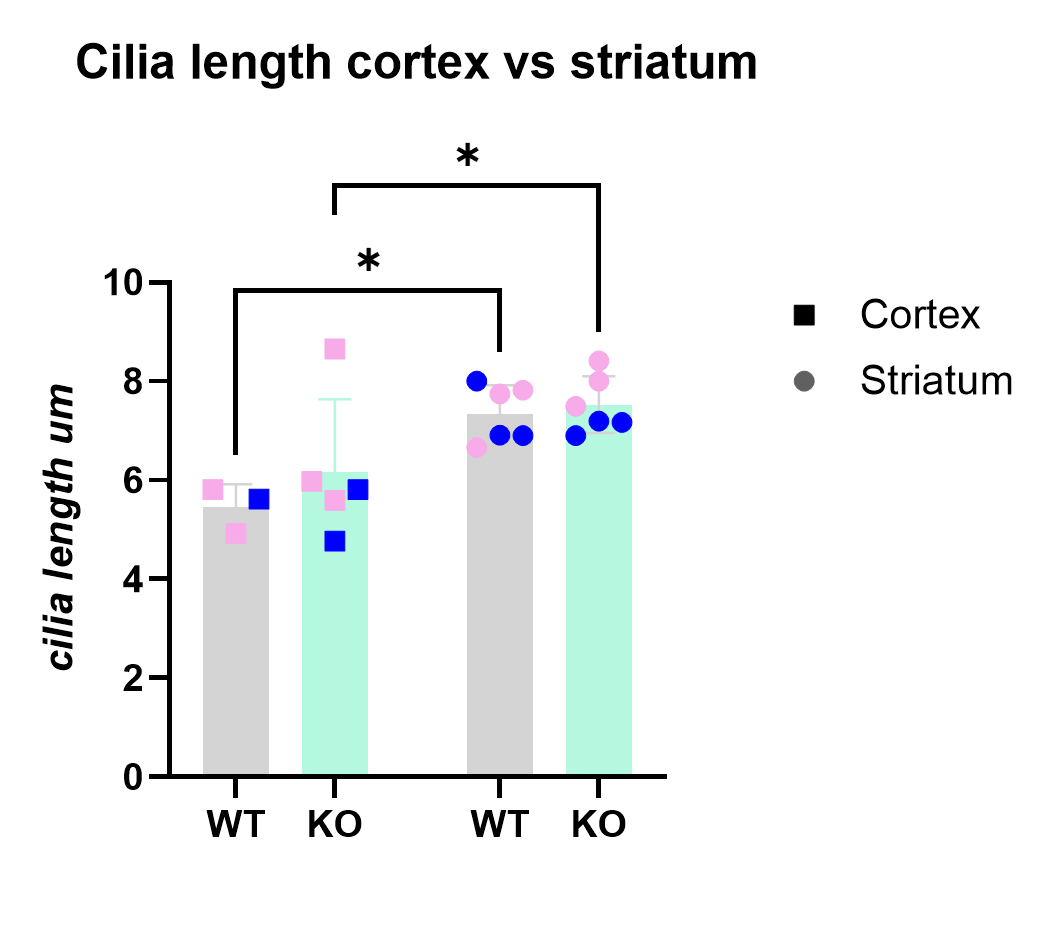

**Figure S3. Quantitative Ai derived data of cilia length (um) between cortex and striatum.** The average of cilia length is shown in µm for both brain regions and genotypes. Statistical analysis was performed using a two-way ANOVA, Tukey’s test. Significant differences were observed between cortex and striatum in cilia length. Blue indicates male, pink indicates female. **P* <0.05

| **Experiment day** | **Antigen** | **Primary** | **Secondary** | **False Positives** | **Figure** |
| --- | --- | --- | --- | --- | --- |
| Day 1 | GPR88-Venus  +  AC3 | Ch anti-Venus  Rb anti-AC3 |  |  |  |
| Day 2 (morning) |  |  | Anti-ch 488  Anti-rb 594 | None |  |
| Day 2 (afternoon) | SATB2  or  DARPP-32 | Rb anti-SATB2  or  Rb anti-DARPP-32 |  |  |  |
| Day 3 | SATB2  or  DARPP-32 |  | Anti-rb 647 | Alexa 647 may detect remaining unbound anti-AC3 and anti-SATB2 or anti-AC3 and anti-DARPP-32 | **Fig 3A, Fig 3E** |

**Table S1:** Experimental timeline of sequential staining for SATB2 and DARPP-32 (**Figure 3A and 3E**).
